# A microfluidic platform for the observation and quantification of fungal highways

**DOI:** 10.1101/2025.09.16.676496

**Authors:** Eleonora Moratto, Emily Masters-Clark, Amelia J. Clark, Aliya Pachmann, Saskia Bindshedler, Pilar Junier, Claire E. Stanley

**Author notes:** These authors have contributed equally (Eleonora Moratto, Emily Masters-Clark).

## Abstract

Soil is a complex system characterised by intra- and inter-kingdom interactions among microbial communities. While many different types of fungal-bacterial interactions have been described, hyphal-mediated transport of bacteria via the so-called “fungal highway” (FH) has not been mechanistically described. Bacteria require a liquid film for active movement; therefore, liquid saturation is a significant limiting factor for their dispersal in soil. Hyphal networks contribute to the connectivity between discrete soil microbial populations by providing a physical network through the unsaturated soil matrix. This network serves as a scaffold for liquid transport, thus allowing bacteria to migrate further or access previously isolated spaces. Studying these interactions is challenging due to the complex and stochastic nature of the soil environment; this “black box” aspect makes it difficult to visualise interactions at the microbial scale. Microfluidic technology can provide a solution by offering precise imaging at a high resolution in a physically and chemically controlled environment. We designed a microfluidic device, the Fungal Highways Device (FHD), that allows us to culture filamentous organisms in unsaturated environments and visualise and quantify bacterial dispersal along hyphal networks at the single-cell level. We showed that *Pythium ultimum* is essential for *Pseudomonas putida* movement across an unsaturated environment, and we identified mycelial biomass and hyphal front length as key factors influencing the bacteria’s movement towards the outlet. We propose that the liquid transport facilitated by *P. ultimum* mycelium influences the FH behaviour during its interaction with *P. putida*.

## Introduction

One of the central challenges in microbial ecology is understanding how microbial diversity is structured and sustained, and how this underpins essential ecosystem functions such as nutrient cycling and carbon storage (Delgado-Barquerizo et al., 2016; Singh, J.S, 2015). Hyphal-mediated microbial dispersal is emerging as a key mechanism influencing ecological processes, including resource access, competition, and gene transfer across diverse environments (Dubey, M., et al., 2021; Erktan & Scheu, 2020). Fungal highways (FHs) is a term used to describe the dispersal of prokaryotes along hyphae belonging to Fungi and fungal-like organisms, such as Oomycetes (Kohlmeier et al., 2005; Jansa et al., 2021). The ability of mycelial networks to occupy high volumes of the environment and connect spatially separated micro-niches means that microorganisms can disperse further than their own motility might otherwise allow and expand the range over which they are functionally active. Mycelial networks are well adapted for growing in heterogeneous substrates, such as soil, where they can bridge liquid unsaturated environments and air-filled pores (Sun et al., 2020). This, coupled with hyphae’s ability to transport liquid (Clark et al., 2024; Allen, 2007), facilitates the movement of other microorganisms, which are limited in their dispersal by the need for a continuous liquid film (Álvarez-Barragán et al., 2023).

Despite our significant understanding of the ecosystem functions of fungal highways, we lack mechanistic insight into the processes that drive these interactions. Current methods used to investigate FHs can be split into three main categories: (i) columns and millifluidic devices (Junier et al., 2021; Simon et al., 2015; Kuhn et al., 2022, Buffi et al., 2023), (ii) agar plate-based methods (Zhang et al., 2018; Xiong et al., 2022) and (iii) microcosms (Jiang et al., 2021; Warmink et al., 2009; Yang et al., 2017). Despite the difference in scale, they are all based on the presence of two compartments: one inoculated with the mycelium and prokaryotes of interest, and one containing the target medium, which are then separated by a barrier that cannot be crossed by the dispersing microorganism alone. Sampling of the target medium is then carried out to assess FH occurrence. While all these methods have proven invaluable in identifying which organisms form FHs and what environmental factors influence FHs, few allow FH observation at a cellular level (Mafla-Endara et al., 2021; Buffi et al., 2023). This mainly stems from the challenging environment in which they inhabit. Soil is physically, chemically and biologically complex as well as opaque, all significant barriers to direct observation (Stanley & van der Heijden, 2017). In most cases, plate-based assays cannot restrict hyphae to a single optical plane, resulting in overlapping hyphae and limited hyphal resolution. Furthermore, agar-based assays lack optical transparency and maintain a continuous hydrophilic, liquid-saturated environment, which can be easily crossed by flagellated prokaryotes without the aid of a fungal scaffold. This makes agar-based assays uninformative for studies on how FHs impact connectivity between discrete soil microbial populations under unsaturated conditions.

In recent years, microfluidic devices (Zhao et al., 2014) have become capable of imitating crucial aspects of soil by creating highly regulated physical and chemical conditions and allowing for precise environmental manipulation (Stanley et al., 2016). Microfluidic devices have been well adapted to biological applications, and many have been developed for the specific study of fungi and their interactions (Richter et al., 2022). They offer an excellent platform for conducting these investigations, as they enable real-time, high-resolution imaging at the microscale. The device geometry can be designed to elicit certain behaviours or interactions, and environmental parameters can be controlled or implemented. Several Lab-on-a-Chip technologies were designed to quantify various aspects of filamentous microbial biology, including hyphal tip force (Sun et al., 2020) or fungal space searching and foraging strategies (Held et al., 2019). Furthermore, interaction chips have also been developed for the study of bacterial-fungal interactions (BFIs) (Stanley et al., 2014) as well as fungal-fungal interactions (FFIs) (Gimeno et al., 2021). The BFI device was designed to accommodate inoculation of fungal hyphae and bacteria through separate inlets and observe their interactions in separate microchannels. This enabled quantification of parameters such as changes in hyphal growth and differentiation in mycelial architecture and revealed that *Bacillus subtilis* NCIB 3610 inhibit *Coprinopsis cinerea* hyphal growth through signal secretion (Stanley et al., 2014). However, the BFI device is not suitable for FH observation because it was designed to accommodate full channel saturation with liquid media. Using the FFI device, Gimeno et al. (2021) showed that *Trichoderma rossicum* and *Fusarium graminearum* hyphae can grow across unsaturated channels and would therefore be able to bridge the gaps between liquid-saturated soil micro-niches. Furthermore, Clark et al. (2024) used the FFI device to observe and quantify liquid transport by *Pythium ultimum* hyphae across unsaturated channels, suggesting that hyphae might play an active role in bridging these gaps, given that a 2–10 μm-thick liquid film surrounding hyphae is necessary for bacterial transport (Jiang et al., 2021).

In this paper, we develop the Fungal Highways Device (FHD), a microfluidic device that allows for long-term observation of bacterial dispersal along hyphae in unsaturated conditions. We then used this device to quantitatively describe *Pseudomonas putida* movement along *Pythium ultimum* hyphae, model organisms for the study of FHs (You et al., 2022; Buffi et al., 2025). We show that the presence of *P. ultimum* is necessary for *P. putida* transport across an unsaturated channel and demonstrate that mycelial biomass and the distance reached by the longest hyphae in the mycelium (hyphal front length) are determining factors in the bacteria’s ability to reach the outlet. We hypothesise that liquid transport by *P. ultimum* mycelium drives the FH behaviour when interacting with *P. putida*.

## Materials and Methods

### Microbial culture conditions

Mycelia from the fungal-like Oomycete *Pythium ultimum* M194 was cultured on potato dextrose agar (PDA) (Thermo Fisher Scientific) at 26 °C. Fungal stock cultures were kept on potato dextrose agar (PDA) at 26 ºC until use. Fresh active cultures were prepared 3 days prior to each device inoculation to ensure actively growing mycelium. Active cultures were prepared by cutting plugs from the apical edge of *P. ultimum* mycelia using a cork borer (diameter = 4 mm) from the growing edge and placing them in the centre of a fresh PDA Petri dish, then incubated at 26 °C for 3 days. Importantly, cultures were grown on thick agar (25 mL media into each 10 x 10 cm Petri dish) to yield tall agar plugs. This increases the longevity of the experiment: maintaining humidity, nutrients throughout the protocol, and also remain intact to the end of the experiment to be sampled.

*Pseudomonas putida* KT2440::gfp were maintained on nutrient agar (NA) (Oxoid) plates with gentamycin selection (50 µg/mL). Bacterial overnight cultures were prepared by adding 5 µl (10 µL/mL) gentamycin or 50 ppm kanamycin to 5 mL nutrient broth (NB) (Oxoid). The cultures were inoculated with the desired bacteria from a single colony and incubated overnight, shaking at 200 rpm and 25 °C.

### Device design and fabrication

A digital 2D photomask containing the device design was generated using AutoCAD. Master mould fabrication was performed using standard photolithography techniques. Briefly, a 9 cm silicon wafer was cleaned with solvents and oxygen plasma, then spin-coated (WS-650MZ Modular, Laurell Technologies Corporation) with SU-8 2010 photoresist (Kayaku Advanced Materials) up to 3000 rpm (ramp 350 rpm/s) for 40 seconds to achieve a height of 10 µm. The coated wafer was soft baked (65 °C for 1 min, 95 °C for 3 min), before patterning using a maskless system (Microlight3D) (85% power, 128.5 s exposure per frame). Following UV exposure, the wafer was baked once more (65 °C for 1 min, 95 °C for 4 min) before uncured SU-8 was removed using developer solution for 2 min. The wafer was then silanised for 2 h under a vacuum using 50 μL of chlorotrimethylsilane (Sigma-Aldrich).

Standard soft lithography was used to cast poly(dimethylsiloxane) (PDMS) and manufacture FHDs. Briefly, PDMS (Sylgard 184 elastomer kit, VWR) was mixed in a 10:1 ratio of base to curing agent and degassed for 1 h in a vacuum chamber to remove bubbles. This was then poured onto the centre of the master mould and cured at 70 °C overnight. Glass-bottomed Petri dishes (World Precision Instruments; Ø dish = 50 mm, Ø glass = 23 mm) were coated with 0.5 g of PDMS using a NiLO spin coater (500 rpm, 25 ms^-1^ acceleration, for 50 s) and cured at 70 °C for 2 h.

The cured PDMS was removed from the mould, cut into individual slabs and the inlets punched using a precision cutter (SLS; diameter = 4.75 mm). The PDMS slabs were then washed: 5 mins sequential sonication in 0.5 mM NaOH, 70% EtOH, and finally sterile Milli-Q water, rinsing with sterile water between each step. The PDMS-coated Petri dishes were also washed, but without the sonication and incubation; these were simply filled with each washing reagent in sequence to minimise any interaction with the plastic outer edge. The PDMS slabs and coated Petri dishes were dried with filtered pressurised air and left at ~1 h at 70 °C. Once dry, they were next placed into a plasma oven and treated with air plasma for 1 min (Diener Electronic, Germany; vacuum pressure 0.75 mbar, power 50%), after which they are removed and immediately bonded together to form the FHD. The fabricated FHD was then placed back into the oven at 70 °C for 48 h to ensure hydrophobic recovery of the PDMS. Devices were cooled to room temperature before inoculation.

### Device inoculation

*P. ultimum* inoculant agar plugs were cut from the growing edge of the cultured mycelium using a sterile 4 mm-diameter borer and placed into the fungal inlet, mycelium side down (0 days post inoculation (dpi)). An uninoculated PDA plug was placed in the outlet respectively and 100 µL of sterile Milli-Q water was added to the dish around the device to maintain a humid environment. The inoculated devices were then incubated at 26 °C *overnight*.

The next day (1 dpi) the desired bacterial culture was centrifuged at 2000 rpm for 10 mins. After centrifugation, the supernatant was discarded, and the pellet resuspended in 5 ml sterile 1X Phosphate-Buffered Saline (PBS) (90% Milli-Q water, 10% Gibco™ PBS, pH 7.2). This washing step was repeated twice to remove the culture media, and the final pellet was resuspended in 3 ml sterile PSB to concentrate bacteria. The optical density at 600 nm (OD600) of this final suspension was read and then adjusted to a final OD of 1.6. 10 µl of this bacterial suspension was pipetted directly onto the top of the pre-cut *P. ultimum* plug in the FHD inlet and allowed to dry. Then, a fresh PDA plug was placed on top in contact with the fungal plug and the device incubated at 26 °C for the remainder of the experiment.

### Imaging and image analysis

Devices were imaged every 24 h (1-6 dpi) using inverted 137 microscopes (Eclipse Ti-U and Eclipse Ti-2, Nikon). The entire FHD channel was imaged by tile scanning at 20x using phase contrast (PC) and green fluorescence microscopy (FITC) filters. Mycelial, bacterial and liquid locations were quantified from the tile scans using an ImageJ(Fiji) (Schindelin et al., 2012) macro developed for this project (Supplementary Macro 1). Briefly, a line matching the FHD channel thickness was drawn along the channel, and manual thresholding was applied to the PC image to distinguish the mycelial structure from the background. The image was then made into a binary image to normalise the values across repeats. The average pixel intensity was measured along the line for PC and FITC. The FITC values were then normalised based on background fluorescence. Hyphal front length was quantified using ImageJ(Fiji) (Schindelin et al., 2012) to measure the distance from the channel inlet to the furthest hyphal tip of the mycelium. The bacterial and hyphal biomasses in the channel were obtained by averaging the normalised pixel intensities of the FITC and PC channels, respectively. All data was analysed in R Studio, using packages ggplot2, dyplr and matrixStats. We also acquired exploratory videos or images at 40x, switching between PC and FITC, using in-built autoLUT software.

### Bacterial transport quantification

To quantify the amount of bacteria that traversed the FHD, we used *in vitro* assessment of colony-forming units (CFUs). A sterile 4 mm borer was placed into the target inlet and twisted to sever the hyphae growing into the target plug from the channel. The target plug was removed and placed in 1 ml sterile-filtered 1% hexametaphosphate to separate the bacteria that may be adhered to the hyphae. Using a pestle or forceps, the agar plug was ground in the water until homogenous. The suspension was pipetted into 9 ml sterile-filtered 1X PBS (90% MilliQ water, 10% Gibco™ PBS, pH 7.2) and vortexed gently. The suspension was filtered twice through 20 µm nylon meshes (Plastok Associated LTD), and once through 5 µm meshes (Plastok Associated LTD). 100 µl of the filtered suspension was pipetted onto an NB agar plate with the appropriate antibiotic and incubated at 27 °C for three days. CFUs were counted by taking pictures of the plates and using the cell counter tool in ImageJ software.

## Results

### Design and function of the Fungal Highways Device

To observe and quantify how bacteria disperse along hyphal networks, we developed the Fungal Highways Device (FHD). This device enables unidirectional growth of hyphae and co-inoculation with bacteria in unsaturated conditions, allowing us to monitor network topology and position relative to one another over several days (Figure 1A-D). To develop and characterise the FHD, we use the FH model organisms *Pythium ultimum* and GFP-tagged *Pseudomonas putida* (You et al., 2022; Buffi et al., 2025).

**Figure 1.**
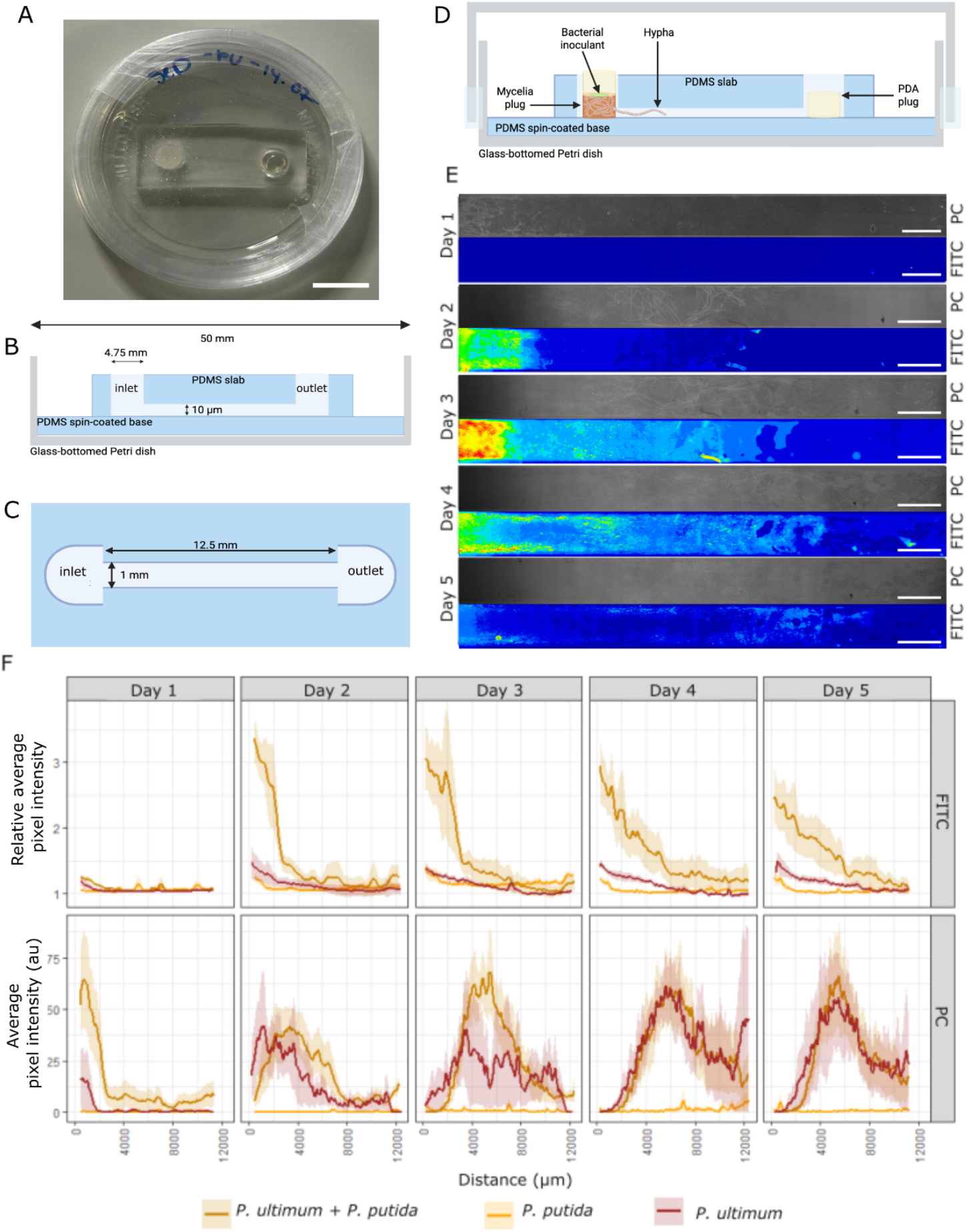
The Fungal Highways Device (FHD) allows visualisation and quantification of bacterial dispersal across hyphal networks under unsaturated conditions. **A)** Photograph of an inoculated FHD showing a poly(dimethylsiloxane) (PDMS) slab with the embossed microchannel bonded to a PDMS-coated glass-bottomed Petri dish. **B)** Schematic representation of the FHD, side view (not to scale - dimensions given). The glass-bottomed Petri dish is spin-coated with a thin layer of PDMS, to which the PDMS slab embossed with 10 µm channel and inlets is bonded. **C)** Schematic representation of the FHD, top view (not to scale - dimensions given), which shows the 2D design of the microchannel with dimensions. **D)** Schematic representation of the FHD inoculated with both mycelia and bacteria, and a hypha moving through the channel from the inlet to the outlet (containing a fresh PDA plug). The inoculation strategy was designed to allow hyphae to pre-establish in the device for one day prior to bacterial co-inoculation. **E)** Representative images of the FHD channel inoculated with *P. ultimum* and *P. putida-*GFP 1 to 5 days post-mycelial-inoculation (dpmi). Bacteria were inoculated on day 1 after imaging. Top panels are acquired using phase contrast (PC) and bottom panels using FITC, enabling the visualisation of the mycelium and bacteria, respectively. The FITC image is displayed with a 16-colour LUT look-up table with blue representing the low pixel intensity and red representing high pixel intensity. Scale bars, 1 mm. **F)** Quantification of the relative average pixel intensity of the FITC and the average pixel intensity for PC images (Supplementary Macro 1) along the channel each day of devices inoculated with *P. ultimum* and *P. putida* (orange), *P. putida* only (yellow) and *P. ultimum* only (red). The solid line represents the mean value and the corresponding shaded area, the standard error (n = 8). Panels B, C & D were created in BioRender. Pachmann, A. (2025) https://BioRender.com/00i19ns

The FHD design consists of an observation channel (width: 1000 µm, length: 12.5 mm) to allow for the formation of a substantial mycelial network, upon which a sufficient hyphal-mediated liquid film can be supported without restricted channel geometry that would disrupt liquid film formation (Figure 1B & C) (Clark et al., 2024). The channel was designed to be 10 µm in height, which has been found to accommodate and confine hyphae on the order of 1-7 μm (Gimeno et al., 2021, Stanley et al., 2014) with *P. ultimum* hyphal diameter ranging between 1.3 to 6.2 μm (Garibaldi et al., 2010). Hyphal or nutrient media plugs can be placed into a 4-mm-diameter inlet or outlet, respectively (located at either end of the main observation channel), allowing for directional growth of the hyphae across the main channel (Figure 1A-C).

The FHD allows for co-inoculation of mycelium and bacteria in the same inlet by first placing a hyphal plug into the inlet and then inoculating 10 µL of bacterial inoculum on top (Figure 1D&E). This is then topped by a second PDA plug, which prevents the bacterial inoculum from drying out as well as providing enough nutrients to sustain the growth of both these microorganisms. Importantly, the design allows for the maintenance of an unsaturated, air-filled channel. Since *P. putida* are flagellated and therefore motile in liquid environments, a fully saturated channel is not suitable for FH visualisation since bacterial motility would act as the dominant form of transport. The unsaturated channel ensures true hyphal-mediated dispersal of bacteria, where liquid is only drawn into the observation channel in the presence of hyphae. To achieve this, the FHD was assembled by bonding the poly(dimethylsiloxane) (PDMS) slab containing the embossed microchannel to a PDMS-coated glass-bottomed Petri dish (Figure 1D). We hypothesised that the hydrophobic nature of PDMS would prevent full liquid saturation of the microchannel, thus preventing bacteria from entering the channel and reducing condensation. To verify this, we quantified the mean water condensation area observed using both phase contrast (PC) and fluorescence (FITC) imaging. We observed that coating of the glass-bottomed Petri dishes with a thin layer of PDMS not only prevented liquid from entering the channel in the absence of hyphae, but also significantly reduced condensation after 48 h (Supplementary Figure 1). To verify if unsaturated conditions prevented bacterial transport, we performed control experiments by inoculating the device with *P. putida* only and imaging the channel with phase contrast (PC) and fluorescence (FITC) microscopy over 5 days. We observed no presence of bacteria in the channel (Figure 1F and Supplementary Figure 2 A&C).

**Figure 2:**
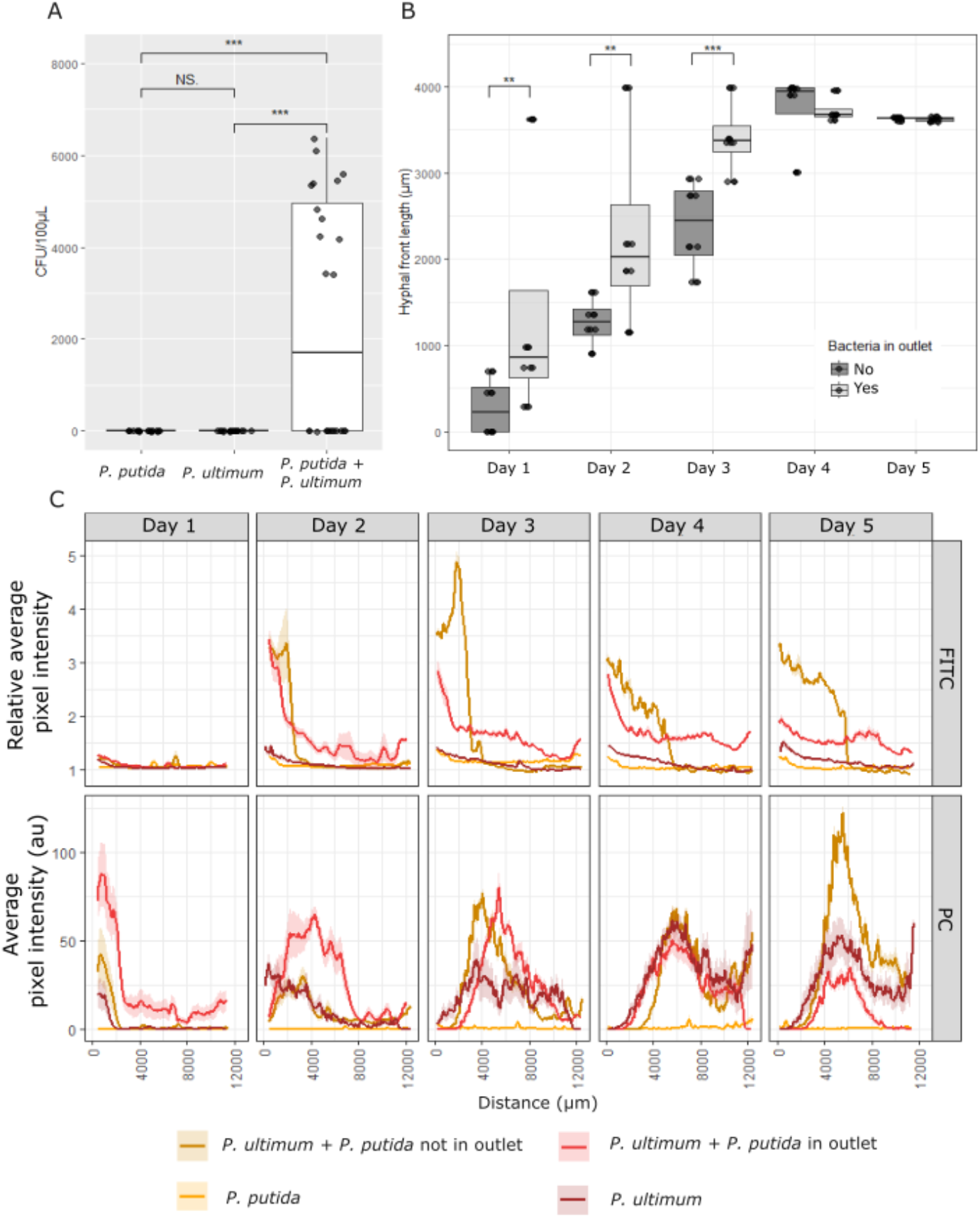
*P. ultimum* hyphal networks enable dispersal of *P. putida* bacteria across a 12 mm unsaturated space in 5 days. **A)** Quantification of *P. putida* colony forming units (CFU/100 µL) extracted from the outlet of FHDs inoculated with (i) *P. putida* only (n = 6), (ii) *P. ultimum* only (n = 6) and (iii) both *P. ultimum* and *P. putida* (n = 8) at 5 dpmi. Three plates were plated per extraction, and each black dot represents the CFU/100 µL of each plate. A Kruskal-Wallis test was used to identify significant differences. Significance is indicated by an asterisk (NS p<0.05; * p ≤ 0.05; ** p ≤ 0.01; *** p ≤ 0.001). **B)** Comparison of hyphal front length from FHDs inoculated with *P. ultimum* and *P. putida* (n = 8) split based on the presence (light grey) or absence (dark grey) of bacteria in the outlet. A Dunn’s post-hoc test was run to identify pairwise differences and significance is indicated by an asterisk (* *p* ≤ 0.05; ** *p* ≤ 0.01; *** *p* ≤ 0.001). **C)** Quantification of the relative average pixel intensity of the FITC and the average pixel intensity for PC images (Supplementary Macro 1) along the channel each day of devices inoculated with (i) *P. ultimum* and *P. putida*, where bacteria were present in the outlet (red, n = 4); (ii) *P. ultimum* and *P. putida*, where bacteria were absent from the outlet (brown, n = 4); (iii) *P. putida* only (yellow, n = 6); and, (iv) *P. ultimum* only (dark red, n = 6). The solid line represents the mean value and the corresponding shaded area, the standard error.

### Developing a quantitative assay to characterise Fungal Highways

To validate the use of the FHD to quantify characteristics of fungal highways such as mycelial network architecture and bacterial dispersal, we developed an experimental pipeline. A *P. ultimum* plug was inoculated in the inlet of a FHD and incubated for 24 h at 26 °C to allow the mycelium to begin colonising the main observation channel. The entire channel was then imaged under PC and FITC filters before inoculating *P. putida* on top of the fungal plug. The device was incubated at 26 °C and imaged every 24 h for 5 days. The images obtained show gradual growth of the mycelium across the channel with hyphal front reaching the second inlet by day 4 (Figure 1E). *P. putida* can be observed moving across the channel and reaching the outlet in 5 days when *P. ultimum* is present, but not in the absence of a mycelium (Supplementary Figure 2A).

To gain further insight into the dynamics of the fungal highway, we developed an ImageJ macro to quantify the average pixel intensity across the channel for each position along the channel (Supplementary Macro 1). The information gathered from the FITC and PC channels provides a proxy for *P. putida* and *P. ultimum*, respectively, i.e., biomass of each organism occupying a certain area along the microchannel (Figure 1F). *P. ultimum* biomass behaves similarly in the presence and absence of *P. putida*, by colonising the channel over time and reaching the second inlet by day 4. As expected, no green fluorescence is detected on day 1, as the bacteria are inoculated directly after imaging *P. ultiumum* on day 1 (Figure 1F). Green fluorescence is only detected in the channel when both organisms are present, indicating that *P. putida* is unable to traverse unsaturated environments (Figure 1F). Notably, most of the bacterial biomass is located behind the hyphal biomass, strengthening our hypothesis that hyphae are driving bacterial movement (Supplementary Figure 2C).

A limitation of this method is the slight hyphal autofluorescence in the FITC channel; however, *P. ultimum* hyphal autofluorescence (1.155 ± 0.0003) is negligible compared to bacterial fluorescence in the presence of *P. ultimum* (1.584 ± 0.001) and similar to the background fluorescence detected when the channel is empty (1.049 ± 0.0002) (Figure 1F, FITC Day 4 & 5 red and brown lines). Another limitation is the macro’s inability to detect hyphae when liquid saturation occurs. For this reason, the majority of *P. ultimum* biomass appears to be concentrated in the middle of the channel between days 3 and 5. This is only due to liquid from the inlet saturating the first half of the channel by day 3 and liquid from the outlet distributing along the mycelium on days 4 to 5 (Figure 1E).

### *Characterisation of* P. putida *dispersal across the FHD channel via* P. ultimum *hyphae*

To ensure the hyphal system could accommodate bacterial dispersal from the inlet to the outlet, we extracted bacteria from the PDA plug and plated them to quantify the colony-forming units (Supplementary Figure 1A). We observed *P. putida* colony formation only in devices inoculated with both *P. ultimum* and the bacteria (Figure 2A), further confirming that hyphae are required for bacterial dispersal in unsaturated conditions. However, we noted that not all co-inoculated devices led to the presence of bacteria in the outlet.

To shed more light on the *P. ultimum* and *P. putida* fungal highway interaction, we split our analysis based on the presence or absence of bacteria in the outlet. We observed that when bacteria were detected in the outlet, the hyphal front had progressed significantly further along the channel during the first three days of growth (Figure 2B & C). Hyphae reaching the end of the channel by day 4 even when bacteria were not detected in the outlet, indicating that hyphal front positioning alone was not sufficient to push the bacteria to the outlet (Figure 2B & C). We note no difference in the relative average green fluorescence intensity of the channel in either case (i.e., regardless of whether or not bacteria were detected in the outlet), indicating that bacterial biomass does not explain the difference in outcome (Figure 3C). The presence of *P. putida* in the channel has no impact on the *P. ultimum* hyphal front or biomass (Figure 3A & B). The difference in hyphal biomass on day 1 can only be attributed to biological variation, since *P. putida* is only added after day 1 imaging. Furthermore, the hyphal surface area is higher on days 1 and 2 when the *P. putida* reaches the outlet (Figure 3C). We also note that when bacteria were not detected in the outlet, the bacterial front does not proceed further than halfway through the channel (Figure 2C). When observing the position of the bacterial front fluorescence peak compared to the hyphal PC peak on day 5, we notice a higher peak in the middle of the device is visible when the bacteria do not reach the outlet (Figure 2C, Supplementary Figure 3). This is supported by the lower *P. ultimum* biomass observed on day 5 (Figure 3C). Liquid saturation in the channel decreases image contrast in the PC channel, rendering the macro less able to detect hyphae. This suggests that when the outlet contains bacteria, the middle of the channel is completely saturated, thus “hiding” some of the hyphal biomass (Supplementary Figure 3). The peak in hyphal biomass detected at the centre of the device on day 5, when bacteria are not found in the outlet (Figure 2C, Supplementary Figure 3), is not an increase in mycelial biomass but a result of unsaturated conditions, making the hyphae in the region better detectable by our macro. Unsaturated conditions remaining in the centre of the device might constitute enough of a barrier to bacterial transport to prevent it from reaching the outlet.

**Figure 3:**
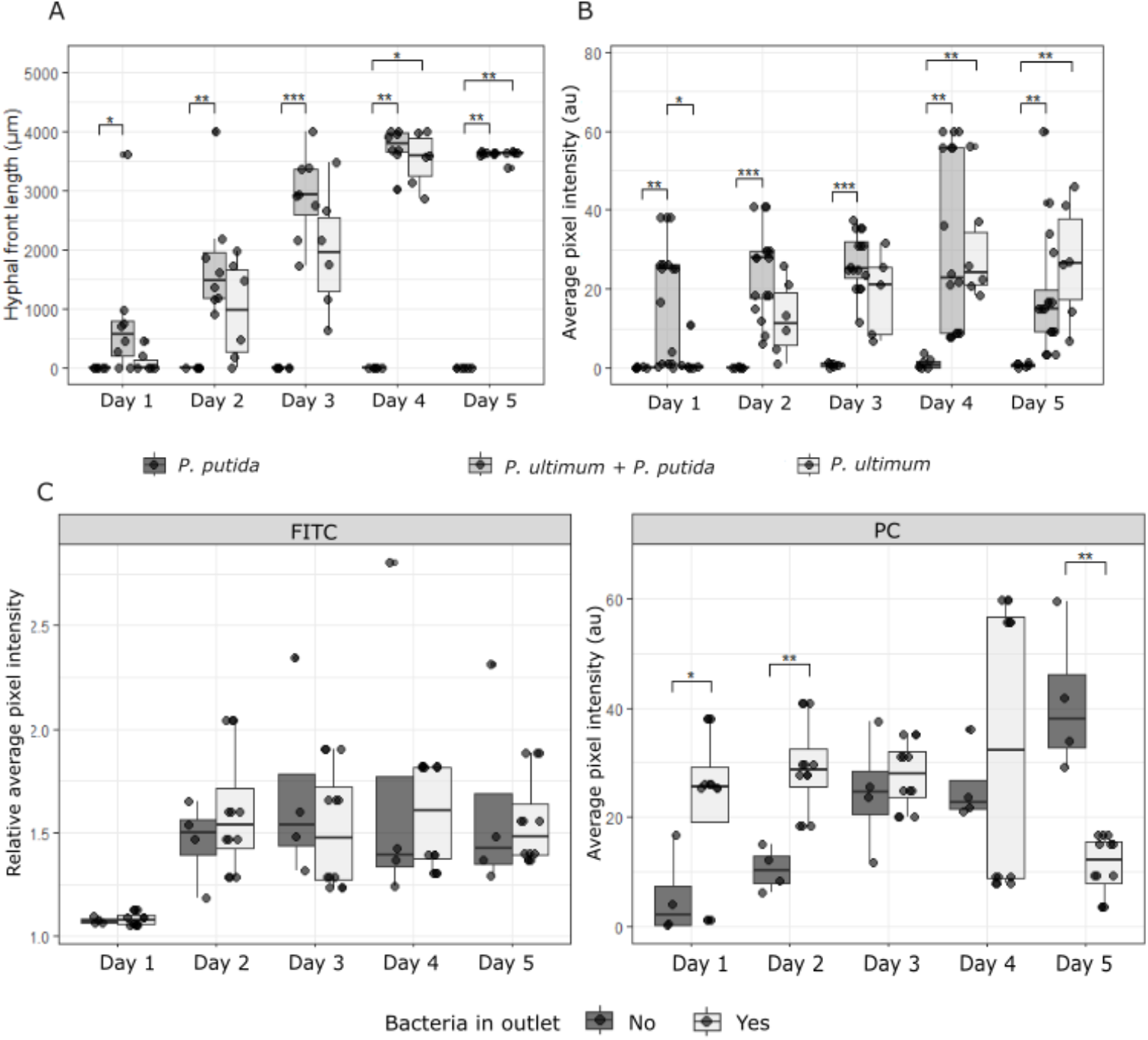
*P. putida* has no impact on *P. ultimum* growth. **A)** Comparison of hyphal front length in FHDs inoculated with (i) *P. putida* only (dark grey, n = 6), (ii) *P. ultimum* only (light grey, n = 6), (iii) both *P. ultimum* and *P. putida* (grey, n = 8) each day. A Dunn’s post-hoc test was run to identify pairwise differences, and significance is indicated by an asterisk (* *p* ≤ 0.05; ** *p* ≤ 0.01; *** *p* ≤ 0.001). **B)** Comparison of average pixel intensity across the channel for FHDs inoculated with (i) *P. putida* only (dark grey, n = 6), (ii) *P. ultimum* only (light grey, n = 6), (iii) both *P. ultimum* and *P. putida* (grey, n = 8) each day. A Dunn’s post-hoc test was run to identify pairwise differences, and significance is indicated by an asterisk (* *p* ≤ 0.05; ** *p* ≤ 0.01; *** *p* ≤ 0.001). **C)** Comparison of FITC relative average pixel intensity and PC average pixel intensity across the channel for FHDs inoculated with both *P. ultimum* and *P. putida* (n = 8), split based on the presence (light grey) or absence (dark grey) of bacteria in the outlet. A Dunn’s post-hoc test was run to identify pairwise differences, and significance is indicated by an asterisk (* *p* ≤ 0.05; ** *p* ≤ 0.01).

### P. putida *uses fungal highways to travel between discrete liquid patches*

To further characterise *P. putida* behaviours moving along *P. ultimum* hyphae and within liquid films, we obtained exploratory real-time videos and images of the hyphal and bacterial front at higher magnification.

We observed that bacteria were never in air-filled gaps, only within liquid films where they accumulate at their perimeter (Figure 4A&B, Supplementary Video 1-5). When discrete liquid patches were present, bacteria were mostly present in contiguous liquid patches (26/26, Table 1) and usually absent from discontiguous ones (18/26, Table 1) (Figure 4A, B, E & F, Supplementary Videos 1,2 & 5). While we occasionally observed bacteria in discontiguous liquid patches, their frequency was lower (Table 1, Figure 4F). We frequently observed active bacterial movement between liquid patches along hyphae (Figure 4A-C, Supplementary Videos 1-3) as well as pulsing liquid films causing rapid movement of bacteria and occasionally inversion of the bacterial flow (Supplementary Video 5) (Table 1). Finally, we observed cytoplasmic streaming in *P. ultimum* hyphae as well as other oomycete structures delimiting different pools of different bacterial concentrations in otherwise contiguous liquid pools (Figure 4D, Supplementary Video 4).

**Table 1:** Summary of observed FH behaviours and their frequency based on a total of 26 observations. Schematic definition of the behaviours can be found in Figure 4F.

| Observed behaviour | Number of observations/26 |
| --- | --- |
| Bacterial presence exclusively inside liquid films | 26/26 |
| Bacterial presence in contiguous liquid patches | 26/26 |
| Bacterial presence in discontinuous liquid patches | 8/26 |
| Bacteria move between liquid pools along hyphae | 20/26 |
| Pulsing of the liquid film | 5/26 |
| Cytoplasmic streaming in hyphae | 9/26 |
| Bacteria trapped in liquid pools by hyphal structures | 8/26 |

**Figure 4:**
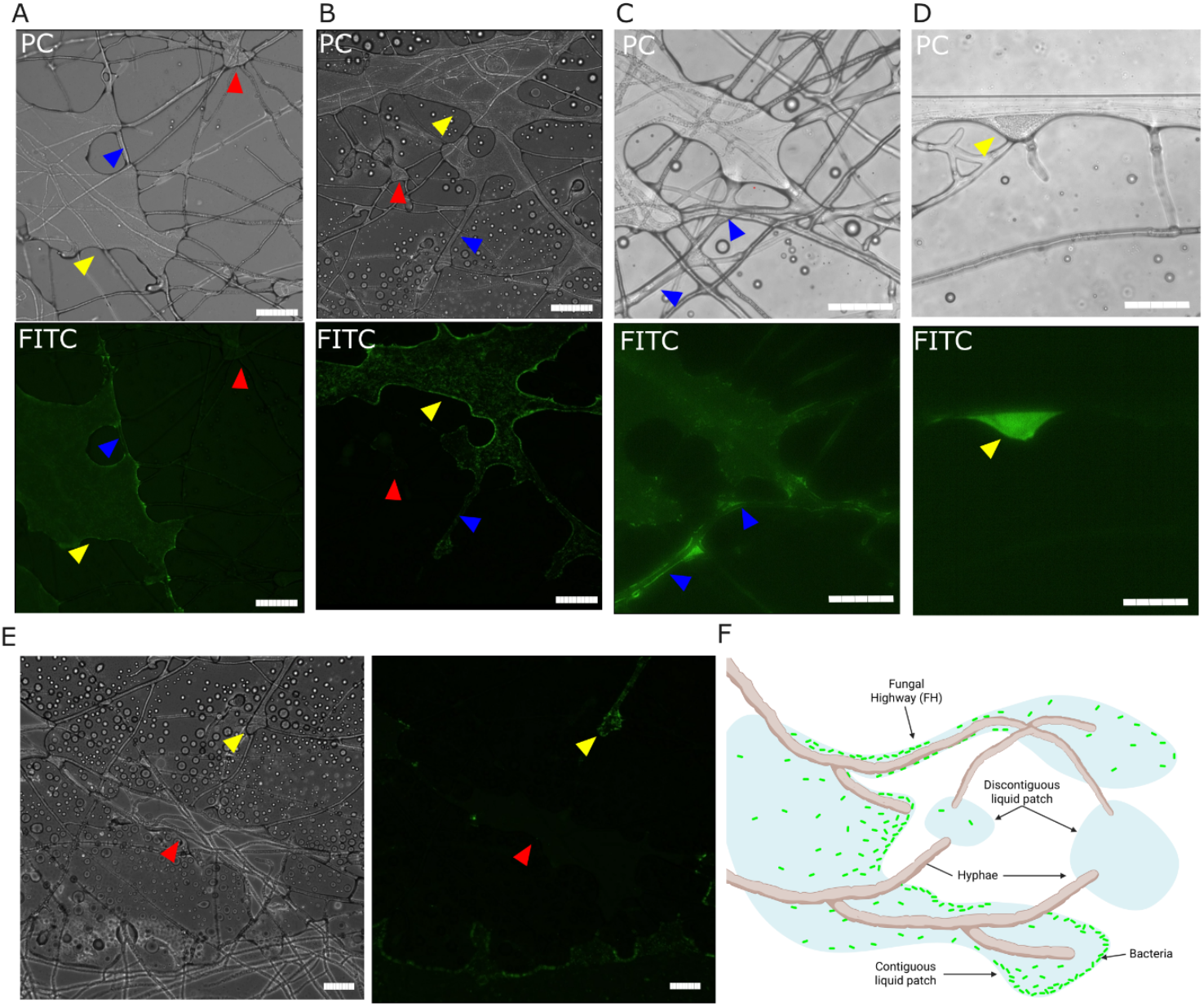
Representative images of FH behaviours. **A), B)** and **C)** *P. putida* bacteria occupy contiguous liquid film patches along hyphae (yellow arrows) and tend to accumulate at the boundaries of the liquid film. Bacteria are absent from most discontiguous liquid films (red arrow). Bacteria move along narrow liquid films along hyphae (blue arrows). Scale bar, 50 µm. **D)** *P. putida* accumulates in liquid films areas delimited *by P. ultimum* structures (yellow arrows). Scale bar, 50 µm. In all cases, the top panel was acquired under phase contrast (PC) and the bottom panel with the FITC filter. **E)** Bacteria are absent from most discontiguous liquid films (red arrow). The left panel was acquired in phase contrast (PC), and the right panel was acquired with the FITC filter. Scale bar, 100 µm. **F)** Schematic representation of the observed behaviours. Bacteria are present in contiguous liquid patches and mostly accumulate at the edge of the liquid. Bacteria invade contiguous liquid patches when connected by hyphae surrounded by a liquid film. In most cases, discontiguous liquid patches are devoid of bacteria. Panel F created in BioRender. Pachmann, A. (2025) https://BioRender.com/j4h01nk

## Discussion

Here, we present a microfluidic platform to quantitatively investigate fungal highway biology at a cellular level. Compared to previously used methods (Junier et al., 2021; Simon et al., 2015; Kuhn et al., 2022; Zhang et al., 2018; Xiong et al., 2022; Jiang et al., 2021; Warmink et al., 2009; Yang et al., 2017), the FHD allows direct observation of the mycelial-bacterial interaction in an unsaturated environment, enabling a variety of resolutions from the organism to the cellular level depending on the magnification used. Furthermore, the FHD can also be used to marry quantitative data obtained through image analysis with information derived from sampling the outlet. In particular, the use of a PDMS-coated base renders the entire channel universally hydrophobic. This provides two important advantages: (i) the reduction in condensation increases image quality, making the FHD ideal for imaging at high magnification, and (ii) hyphal-mediated liquid transport generates more discreet patches akin to a soil environment. This hydrophobic and unsaturated nature of the FHD is essential for investigating true FH behaviour by ensuring authentic hyphal-mediated dispersal of bacteria, particularly when the bacteria of interest are flagellated and can navigate a saturated environment independently. In contrast to other microfluidic devices designed to investigate microorganism interactions (Sun et al., 2020; Held et al., 2019; Stanley et al., 2014; Gimeno et al., 2021), the wide channel design of the FHD and absence of constriction points allows ample space for mycelial growth, liquid and bacterial movement. The presence of an open outlet that allows for the sampling of the target media is crucial for providing both confirmation of bacterial transport, as we presented in this paper, but, analogously to columns and millifluidic methods (Junier et al., 2021; Simon et al., 2015; Kuhn et al., 2022), for the analysis of other parameters such as chemical changes in the environment or gene expression changes in the microorganisms studied.

By coupling the FHD with image analysis, we were able to obtain a precise description of the *P. ultimum-P. putida* interaction dynamics. We confirmed that *P. ultimum* is required for transport of *P. putida* across unsaturated environments (You et al., 2022; Buffi et al., 2025) and only observed the presence of bacteria within liquid patches. We also showed that the presence of air gaps interrupting the liquid film in the centre of the channel was a sufficient barrier for *P. putida*, which accumulated at the edge of the air gaps and did not reach the outlet. When liquid patches were not contiguous or connected by hyphae surrounded by thin liquid films, we observed the absence of bacteria from said patches. Furthermore, discontiguous patches connected by very few hyphae had a lower bacteria concentration. While better qualification of this observational data is needed, this suggests that *P. putida* travels along *P. ultimum* externally, rendering it more likely that this interaction is a true hyphal highway and rather than a hyphal subway (Simon et al., 2017).

*P. putida* is a flagellated bacterium and has been shown to mobilise short distances along the water film surrounding hydrophobic textiles such as polyester (Sherry et al., 2023). This might point towards a passive interaction (Kohlmeier et al., 2005), where *P. putida* will occupy any liquid space, regardless of the presence of *P. ultimum*. This could be confirmed by the absence of an effect of *P. putida* on *P. ultimum* growth. If the interaction were an active one, we could expect the presence of *P. putida* to promote mycelial growth to guarantee transport of more liquid to further locations, but we observe that the higher hyphal biomass that allows bacteria to reach the outlet occurs only 50% of the time, suggesting this is a random phenomenon. Despite this, chemical signalling might be present between the two organisms that alters secretion of compounds by the hyphal network that modify liquid transport, such as hydrophobins and other biosurfactants (Allen et a., 2007; Wosten et al., 2015; Silva et al., 2021). While *P. ultimum* and other fungi have been shown to transport liquid (Clark et al., 2024), the mechanism remains unclear. Different mycelial organisms possess different properties in their hyphal extracellular matrix that renders their surface hydrophilic or hydrophobic and are capable of modifying them through the production of hydrophobins and other biosurfactants (Allen et a., 2007; Wosten et al., 2015; Silva et al., 2021). While this is likely not the only factor determining liquid transport (Clark et al., 2024), bacterial species have shown preferences for mycelia with different surface properties. For example, various isolates of *Pseudomonas* sp. crossed a hydrophilic surface, using *Alternaria destruens*, but crossed a hydrophobic surface, using *Fusarium pseudonygamai* (Álvarez-Barragán et al., 2023). *P. putida*, however, has only been found to travel along hydrophilic hyphae (Wick et al., 2007). It is not unreasonable to suggest that specific bacterial species could engage in chemical signalling with mycelia to alter their surface chemistry and thus their capability to transfer liquid.

Overall, we successfully developed a new tool for the direct observation and quantitative study of fungal highways. While a lot remains to be uncovered, the FHD will be an invaluable tool to the advancement of the field. Future studies will utilise it to shed light on the mechanistic interactions between a variety of mycelial species and their micro-interactors. The use of fluorescently tagged species as well as fluorescent markers for liquid will allow better quantification of mycelial growth and liquid transport dynamics (Clark et al., 2024). Alternatively, fluorescent dyes can be used to investigate interaction with ecologically relevant soil microorganisms that are not amenable to transformation (Richter et al., 2023). Furthermore, the ability to sample from both the inlet and the outlet will allow further investigation of chemical and gene expression changes in the system as well as assessing network selectivity of bacterial species and more complex hyphal networks comprising multiple fungal species.

## Acknowledgments

We acknowledge financial support from the Department of Bioengineering at Imperial College London and The Leverhulme Trust (Research Grant Reference: RPG-2020-352).

## References

Allen MF. Mycorrhizal fungi: highways for water and nutrients in arid soils. Vadose zone journal. 2007 May 1;6(2):291–7.

Álvarez-Barragán J, Cravo-Laureau C, Xiong B, Wick LY, Duran R. Marine Fungi Select and Transport Aerobic and Anaerobic Bacterial Populations from Polycyclic Aromatic Hydrocarbon-Contaminated Sediments. mBio. 2023;14:e02761–22.

Buffi M, Cailleau G, Kuhn T, Li Richter X, Stanley CE, Wick LY, Chain PS, Bindschedler S, Junier P, Fungal drops: a novel approach for macro- and microscopic analyses of fungal mycelial growth, microLife, Volume 4, 2023, qad042.

Buffi M, Kuhn T, Gonzalez D, Bindschedler S, Chain PS, Richter XY, Junier P. Assessing the speed of individual bacteria dispersing on mycelial networks. Evolutionary Ecology. 2025 Jan 25:1–3.

Delgado-Baquerizo M, Maestre FT, Reich PB, Jeffries TC, Gaitan JJ, Encinar D, et al. Microbial diversity drives multifunctionality in terrestrial ecosystems. Nat Commun. 2016;7:10541.

Dubey, M., et al., Environmental connectivity controls diversity in soil microbial communities. Communications Biology, 2021. 4(1): p. 492.

Erktan, A., D. Or, and S. Scheu, The physical structure of soil: Determinant and consequence of trophic interactions. Soil Biology and Biochemistry, 2020. 148: p. 107876.

Garibaldi A, Gilardi G, Gullino ML. First Report of Collar and Root Rot Caused by Pythium ultimum on Coriander in Italy. Plant Disease. 2010 Sep;94(9):1167.

Gimeno A, Stanley CE, Ngamenie Z, Hsung MH, Walder F, Schmieder SS, Bindschedler S, Junier P, Keller B, Vogelgsang S. A versatile microfluidic platform measures hyphal interactions between Fusarium graminearum and Clonostachys rosea in real-time. Communications Biology. 2021 Feb 26;4(1):262.

Held M, Kašpar O, Edwards C, Nicolau DV. Intracellular mechanisms of fungal space searching in microenvironments. Proceedings of the National Academy of Sciences. 2019 Jul 2;116(27):13543–52.

Jansa J, Hodge A. Swimming, gliding, or hyphal riding? On microbial migration along the arbuscular mycorrhizal hyphal highway and functional consequences thereof. The New Phytologist. 2021 Apr 1;230(1):14–6.

Jiang, F., Zhang L., Zhou J., George T.S., and Feng G., Arbuscular mycorrhizal fungi enhance mineralisation of organic phosphorus by carrying bacteria along their extraradical hyphae. New Phytologist, 2021. 230(1): p. 304–315.

Junier, P., et al., Democratization of fungal highway columns as a tool to investigate bacteria associated with soil fungi. FEMS Microbiology Ecology, 2021. 97(2).

Kohlmeier S, Smits THM, Ford RM, Keel C, Harms H, Wick LY. Taking the Fungal Highway: Mobilization of Pollutant-Degrading Bacteria by Fungi. Environ Sci Technol. 2005;39:4640–6.

Mafla-Endara, P. M., Arellano-Caicedo, C., Aleklett, K., Pucetaite, M., Ohlsson, P., and Hammer, E. C., Microfluidic chips provide visual access to in situ soil ecology. Communications Biology, 2021. 4(1), 889.

Richter F, Bindschedler S, Calonne-Salmon M, Declerck S, Junier P, Stanley CE. Fungi-on-a-Chip: microfluidic platforms for single-cell studies on fungi. FEMS Microbiology Reviews. 2022 Nov;46(6):fuac039.

Richter, I., Wein, P., Uzum, Z., Stanley, C. E., Krabbe, J., Molloy, E. M., Ferling, I., Hillmann, F. & Hertweck, C. (2023). Transcription activator-like effector protects bacterial endosymbionts from entrapment within fungal hyphae. Current Biology, 33(13), 2646–2656.

Sherry, A., B.M. Dell’Agnese, and J. Scott, Biohybrids: Textile fibres provide scaffolds and highways for microbial translocation. Frontiers in Bioengineering and Biotechnology, 2023. 11.

Schindelin J., Arganda-Carreras I., Frise E., Kaynig V., Longair M., Pietzsch T., Preibisch S., Rueden C., Saalfeld S. and Schmid B., Nature Methods, 2012, 9(676–682).

Simon, A., et al., Exploiting the fungal highway: development of a novel tool for the in situ isolation of bacteria migrating along fungal mycelium. FEMS Microbiology Ecology, 2015. 91(11).

Simon A, Hervé V, Al-Dourobi A, Verrecchia E, Junier P. An in situ inventory of fungi and their associated migrating bacteria in forest soils using fungal highway columns. FEMS Microbiol Ecol. 2017;93:fiw217

Singh, J.S., Microbes Play Major Roles in the Ecosystem Services. Climate Change and Environmental Sustainability, 2015. 3: p. 163.

Stanley CE, Grossmann G, i Solvas XC, deMello AJ. Soil-on-a-Chip: microfluidic platforms for environmental organismal studies. Lab on a Chip. 2016;16(2):228–41.

Stanley CE, Stöckli M, van Swaay D, Sabotič J, Kallio PT, Künzler M, deMello AJ, Aebi M. Probing bacterial–fungal interactions at the single cell level. Integrative Biology. 2014 Oct 22;6(10):935–45.

Stanley CE, van der Heijden MG. Microbiome-on-a-chip: new frontiers in plant–microbiota research. Trends in microbiology. 2017 Aug 1;25(8):610–3.

Sun B, Chen X, Zhang X, Liang A, Whalen JK, McLaughlin NB. Greater fungal and bacterial biomass in soil large macropores under no-tillage than mouldboard ploughing. European Journal of Soil Biology. 2020 Mar 1;97:103155.

Sun Y, Tayagui A, Garrill A, Nock V. Microfluidic platform for integrated compartmentalization of single zoospores, germination and measurement of protrusive force generated by germ tubes. Lab on a Chip. 2020;20(22):4141–51.

Kuhn, T., et al., Design and construction of 3D printed devices to investigate active and passive bacterial dispersal on hydrated surfaces. BMC Biology, 2022. 20(1): p. 203.

Warmink, J.A., R. Nazir, and J.D. Van Elsas, Universal and species-specific bacterial ‘fungiphiles’ in the mycospheres of different basidiomycetous fungi. Environmental Microbiology, 2009. 11(2): p. 300–312.

Wick, L.Y., et al., Effect of Fungal Hyphae on the Access of Bacteria to Phenanthrene in Soil. Environmental Science & Technology, 2007. 41(2): p. 500–505.

Xiong, B.-J., et al., Impact of Fungal Hyphae on Growth and Dispersal of Obligate Anaerobic Bacteria in Aerated Habitats. mBio, 2022. 13(3): p. e00769–22.

Yang, P., M. Zhang, and J.D. van Elsas, Role of flagella and type four pili in the co-migration of Burkholderia terrae BS001 with fungal hyphae through soil. Scientific Reports, 2017. 7(1): p. 2997.

You X, Kallies R, Kühn I, Schmidt M, Harms H, Chatzinotas A, Wick LY. Phage co-transport with hyphal-riding bacteria fuels bacterial invasion in a water-unsaturated microbial model system. The ISME Journal. 2022 May;16(5):1275–83.

Zhang, Y., et al., Fungal networks shape dynamics of bacterial dispersal and community assembly in cheese rind microbiomes. Nature Communications, 2018. 9(1): p. 336.

Zhao, C., Zhong, G., Kim, D.E., Liu, J. and Liu, X., 2014. A portable lab-on-a-chip system for gold-nanoparticle-based colorimetric detection of metal ions in water. Biomicrofluidics, 8(5).

