## Supplementary Material for "A microfluidic platform for the observation and quantification of fungal highways"

* These authors have contributed equally

### Supplementary material

#### *Supplementary figures*

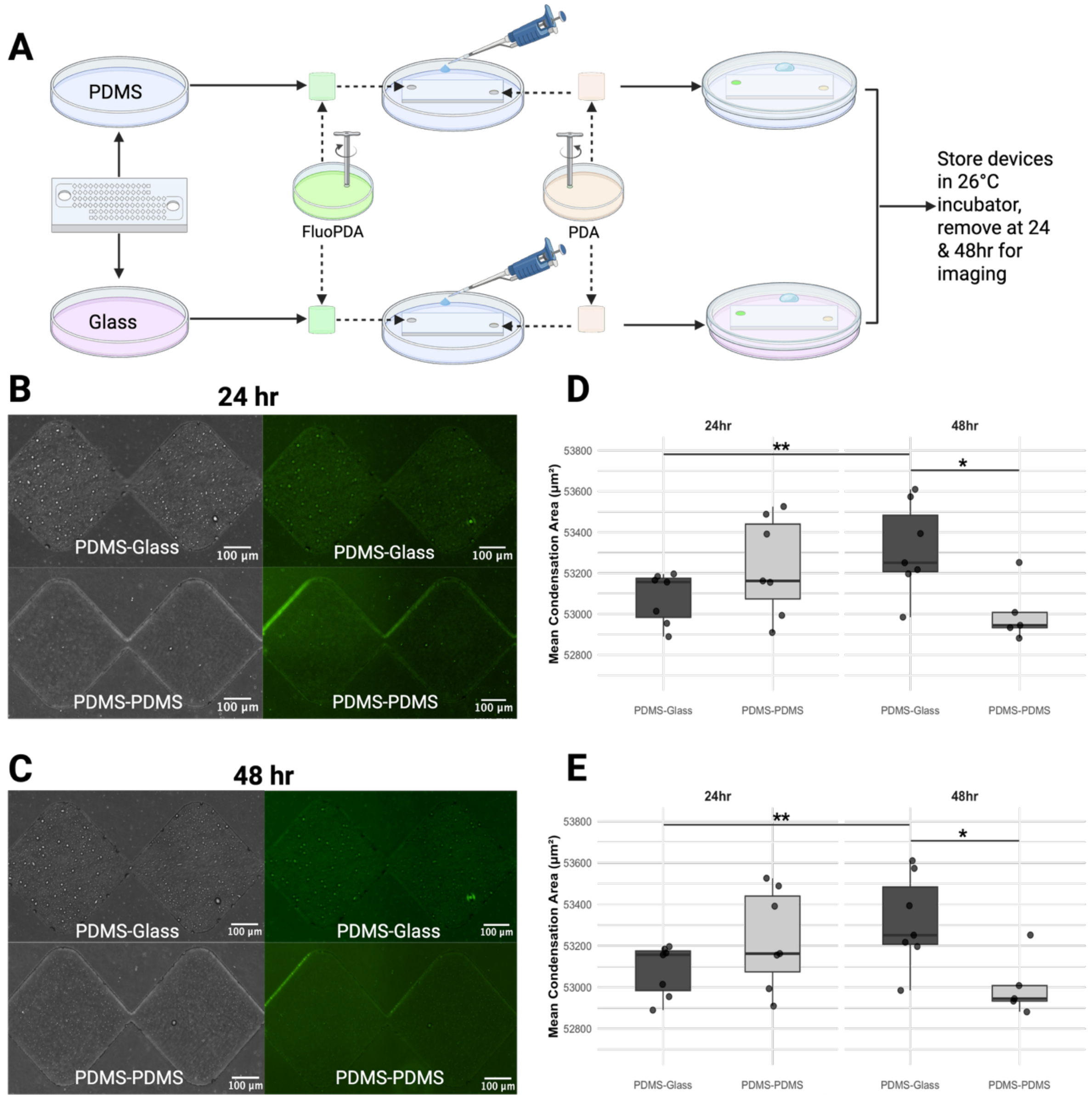


**Supplementary Figure 1: PDMS spin-coating of glass petri dishes decreases condensation.** Panel A illustrates the protocol for investigating the effect of bonding PDMS slabs to either a PDMS or glass base on condensation. To summarise, the cut and washed FFI PDMS slab is bonded to either a glass-bottomed Petri dish with or without a spin-coated layer of PDMS. Then, two agar plugs are placed into the inlets – one plug of FluoPDA (PDA with 0.66 μM fluorescein) and one plug of non-fluorescent PDA. 100 μl of Milli-Q water is added to the petri dish before sealing it with parafilm and storing the devices in an incubator at 26 °C, removing them at 24- and 48-hours post-inoculation for imaging. In the second interaction channel of the FFI device, images of Diamonds 1 and 2 bonded to either PDMS or glass were captured at 24 hours (B) and 48 hours (C) using phase contrast and fluorescence microscopy. Boxplots of the mean condensation area (μm$^2$) of PDMS-glass and PDMS-PDMS devices calculated using phase contrast (D) and fluorescence (E) images showed nearly identical results. Statistical analysis on both phase contrast and fluorescence microscopy datasets yielded the same results. The mean condensation area was analysed using the non-parametric Brunner-Munzel test, appropriate for small, unequal, and heterodaisic datasets. Results showed comparable performance between the setups at 24 hours ($p = 0.4375$), but at 48 hours, the PDMS-glass devices demonstrated significantly more condensation than PDMS-PDMS devices (p = 0.02785). Notably, PDMS-glass exhibited a substantial rise in condensation area from 24 to 48 hours (p = 0.00732), while PDMS-PDMS showed no significant change (p = 0.0949). Created in BioRender. Pachmann, A. (2025) https://BioRender.com/i2notve.

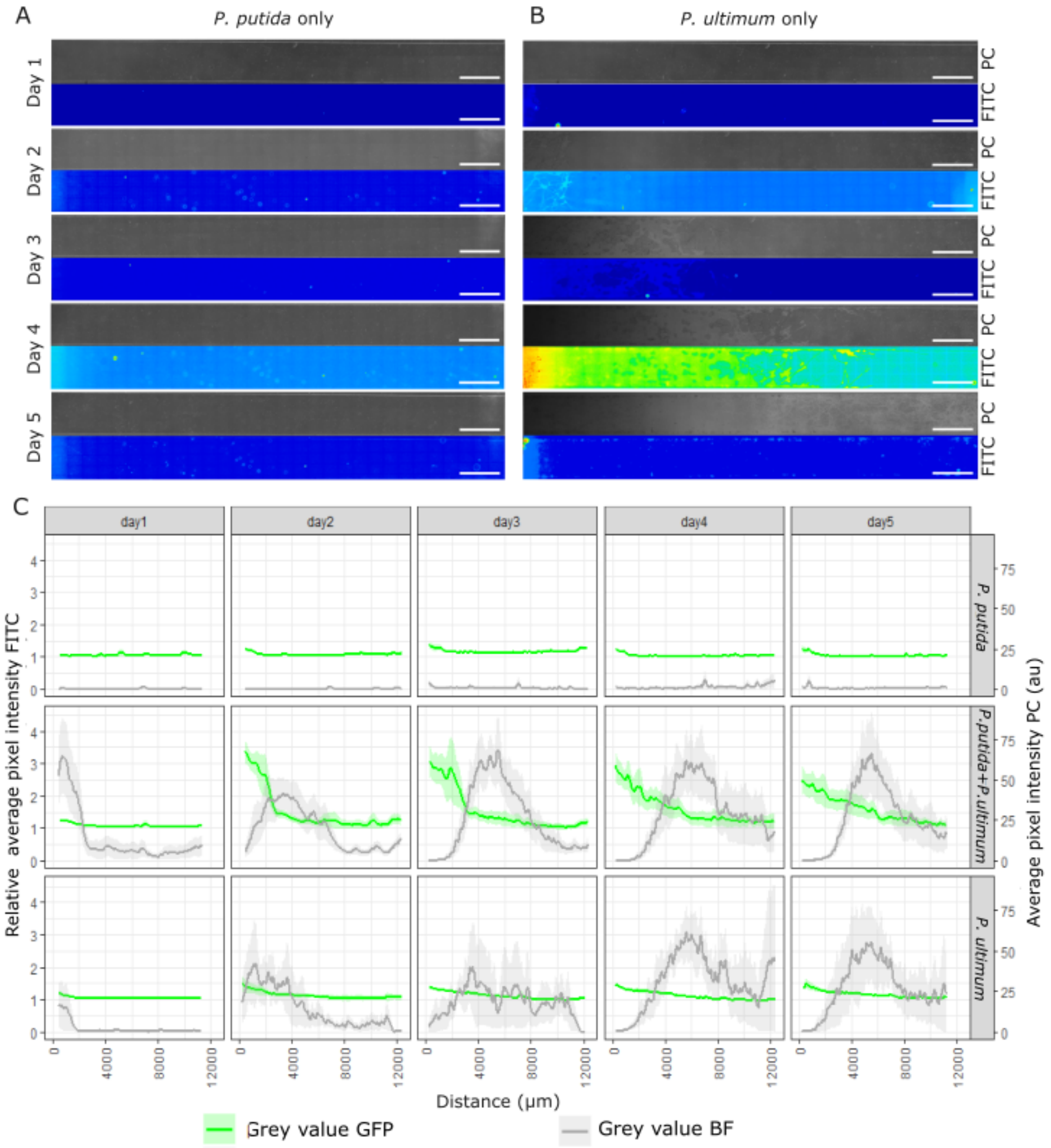


**Supplementary Figure 2: A) & B)** Representative images of the FHD channel inoculated with *P. putida* only (A) and *P. ultimum* only (B) 1 to 5 dpmi. Bacteria were inoculated on day 1 after imaging. Top panels were acquired using phase contrast (PC) and the bottom panels using FITC filter, enabling the visualisation of the mycelium and bacteria, respectively. Scale bars, 1 mm. **C)** Quantification of the relative average pixel intensity of the FITC images (green line) and average pixel intensity of the PC images (grey line) along the FHD channel (see Supplementary Macro 1 for details**).** Images are taken each day of devices inoculated with *P. ultimum* and *P. putida*, *P. putida* only and *P. ultimum* only. The solid line represents the mean value, and the corresponding shaded area, the standard error.

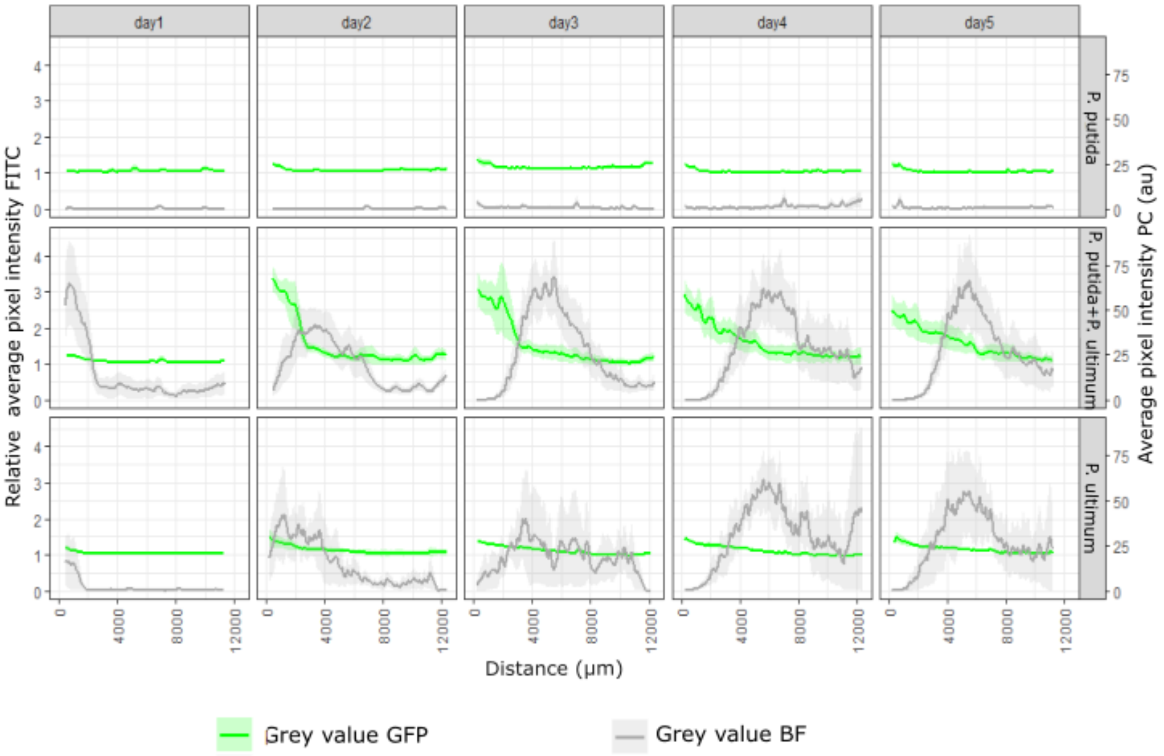


**Supplementary Figure 3:** Quantification of the relative average pixel intensity of the FITC images (green line) and average pixel intensity of the PC images (grey line) along the FHD channel (see Supplementary Macro 1 for details). Images are taken each day of devices inoculated with *P. ultimum* and *P. putida* where bacteria were present in the outlet (n = 4), *P. ultimum* and *P. putida* where bacteria were absent from the outlet (n = 4), *P. putida* only (n = 6), and *P. ultimum* only (n = 6). The solid line represents the mean value, and the corresponding shaded area, the standard error.

### Supplementary Videos

**Supplementary Video 1:** Real-time video of the FHD channel inoculated with *Pythium ultimum* and *Pseudomonas putida*-GFP. The video was recorded in real time at 40x magnification using phase contrast microscopy first and switching to the FITC filter thereafter. *P. putida* swims within liquid patches formed around *P. ultimum* hyphae. The bacteria concentrate at the perimeter of the liquid patch and move along a single hypha surrounded by a fine liquid film.

**Supplementary Video 2:** Real-time video of the FHD channel inoculated with *Pythium ultimum* and *Pseudomonas putida*-GFP. The video was recorded in real time at 40x magnification using phase contrast microscopy first and switching to the FITC filter thereafter. *P. putida* moves within liquid patches along *P. ultimum* hyphae. Bacteria is absent from discontiguous liquid patches.

**Supplementary Video 3:** Real-time video of the FHD channel inoculated with *Pythium ultimum* and *Pseudomonas putida*-GFP. The video was recorded in real time at 40x magnification using phase contrast microscopy first and switching to the FITC filter thereafter. *P. putida* actively moves between contiguous liquid patches along hyphae and pulsing liquid films.

**Supplementary Video 4:** Real-time video of the FHD channel inoculated with *Pythium ultimum* and *Pseudomonas putida*-GFP. The video was recorded in real time at 40x magnification using phase contrast microscopy first and switching to the FITC filter thereafter. High concentration of *P. putida* being trapped by hyphae within a small section of liquid film.

**Supplementary Video 5:** Real-time video of the FHD channel inoculated with *Pythium ultimum* and *Pseudomonas putida*-GFP. The video was recorded in real time at 40x magnification using phase contrast microscopy first and switching to the FITC filter thereafter. Pulsing liquid films cause rapid movement of bacteria and inversion of the bacterial flow along *P. ultimum* hyphae.

### Supplementary Methods

#### *Testing the effect of a PDMS base on condensation*

To test the ability of condensation to form on different surfaces, we created FFI devices (Gimeno et al., 2021) by bonding PDMS slabs to glass-bottomed Petri dishes either with or without a PDMS coating (see *Device design and fabrication* and Supplementary Figure 1A). To observe condensation dynamics independent of biotic factors, methods were adapted from Clark et al.'s protocols for visualising liquid redistribution within the same device (Clark et al., 2024). FFI devices were inoculated with (i) a PDA plug and (ii) a PDA plug supplemented with 0.66 μM filter-sterilised fluorescein (Clark et al., 2024) before being placed in an incubator at 26 °C. Devices were removed at 24 hours post-inoculation (hpi) and 48 hpi for phase contrast (PC) and fluorescence microscopy imaging using a Nikon Eclipse Ti-2 inverted microscope. Microscope images were analysed in FIJI (Schindelin et al., 2012). Images were converted to 8-bit .tiff files, before performing Renyi's entropy-based thresholding and condensation area within the channels was measured and recorded (Supplementary Macros 2 & 3). Statistical analysis was performed in RStudio using the Dplyr, ggplot2, and brunnermunzel packages. The non-parametric Brunner Munzel test was used for significance testing between timepoints and device conditions.

### Supplementary Macros

Here, we report all macros used to conduct semiautomated image analysis within the software ImageJ. All are designed to be opened in ImageJ and run in their entirety, but do require user intervention, which is indicated by instructions appearing on screen. Macro 1 is written in ImageJ Macro language (IJM). The macro is designed to analyse a raw .nd2 file containing the tiled phase contrast image in channel 1 and a fluorescent FITC image in channel 2. The macro outputs three .csv files containing the raw average pixel intensity measured along the device in each channel and a background value for FITC, respectively.

Macros 2 and 3 are written in Jython, a Java implementation of Python. Microscopy files are pre-processed into .tiff hyperstacks with two channels; phase contrast and FITC, and saved into a folder for batch processing with Macro 2 and 3. Macro 2 and Macro 3 are executed in sequential order. Macro 2 is designed to analyse .tiff hyperstacks containing two channels: phase contrast and FITC. It uses user-drawn lines to correct any skew in the image and employs a user-defined bounding box to geometrically transform the region of interest (ROI) pattern onto the image, outlining the channels. The transformed ROI patterns and the deskewed images are then saved into their respective folders as .zip files (for the ROI) and .tiff files (for the two channels). Macro C requires the user to select a folder containing the deskewed images of the channel of interest, along with the .zip files of the transformed ROIs. After selection, the macro thresholds the images using Renyi's entropy thresholding method. It then calculates the total area of each ROI, the area within the ROI that registers above the threshold, and the percentage of the ROI which presents a signal (% signal coverage). Note: this macro requires .tiff microscopy files with scale information stored in their metadata.

***Supplementary Macro 1: FHD image analysis***

```
//Set line width to channel width
run("Set Measurements...", "area mean min perimeter redirect=None decimal=3");
run("Line Width...", "line=3000");
setTool("line");

//Get directory and Image name
dir1 = getDirectory("image");
imageName = getTitle();
// Remove the file extension from the image name
dotIndex = indexOf(imageName, ".");
if (dotIndex != -1) {
   imageName = substring(imageName, 0, dotIndex);
}

//Duplicate image
run("Duplicate...", "duplicate");
rename("Image");

//Draw line
waitForUser("Draw line along the channel, then hit OK");
roiManager("Add");

//Split channels
run("Split Channels");

//Measure line green
selectImage("C2-Image");
roiManager("Select", 0);
run("Plot Profile");
```

```
Plot.getValues(xValues, yValues);
csvFilePath = dir1 + imageName + "_bacteria.csv";
file = File.open(csvFilePath);
print(file, "Distance_microns,Gray_Value");
for (i = 0; i < xValues.length; i++) {
    print(file, xValues[i] + "," + yValues[i]);
}
File.close(file);
//print("Plot profile data saved as: " + csvFilePath);
selectImage("Plot of C2-Image");
run("Close");

//Make hyphae into binary
selectImage("C1-Image");
run("Gaussian Blur...", "sigma=0.7 stack");
run("8-bit");
run("Sharpen");
run("Subtract Background...", "rolling=30");
waitForUser("adjust threshold", "test the threshold parameters in the window that next appears.");
run("Threshold...");
waitForUser("Click Apply then click OK");
run("Dilate");
run("Analyze Particles...", "size=100-Infinity pixel circularity=0.00-0.15 show=Masks");

//Measure line hyphae
selectWindow("Mask of C1-Image");
roiManager("Select", 0);
run("Plot Profile");
Plot.getValues(xValues, yValues);
csvFilePathf = dir1 + imageName + "_fungi.csv";
```

```
filef = File.open(csvFilePathf);
print(filef, "Distance_microns,Gray_Value");
for (i = 0; i < xValues.length; i++) {
   print(filef, xValues[i] + "," + yValues[i]);
}
File.close(filef);
//waitForUser("Save plot data, then press ok");
selectImage("Plot of Mask of C1-Image");
run("Close");

//Measure green background
selectImage("C2-Image");
setTool("rectangle");
waitForUser("Draw Rectangle for background");
roiManager("Add");
roiManager("Select", 1);
roiManager("Measure");
saveAs("Results", dir1+imageName+"_bacteria_n.csv");

//Close everything
roiManager("Delete");
roiManager("Delete");
selectWindow("Results");
run("Close");
selectImage("Mask of C1-Image");
run("Close");
selectImage("C2-Image");
run("Close");
selectImage("C1-Image");
run("Close");
```

***Supplementary Macro 2: Image processing & ROI alignment***

```
# Macro 2 (Jython): Interactive Processing, ROI Alignment, Channel Splitting/Merging


from ij import IJ, ImagePlus
from ij.plugin.frame import RoiManager
from ij.gui import WaitForUserDialog, Roi, PolygonRoi, Line
import os, math
from ij.plugin import ChannelSplitter, RGBStackMerge
from ij.io import FileSaver, DirectoryChooser, OpenDialog


# =======================
# Helper Functions
# =======================


def transform_single_roi(roi, scale_x, scale_y, translate_x, translate_y):
    """
    Transform one ROI by scaling & translating. Adapt to correct type:
    - POLYGON, FREEROI, TRACED_ROI, POLYLINE, FREELINE, ANGLE => PolygonRoi
    - LINE => new Line
    - RECTANGLE => new rectangular Roi
    - Others => skip (return None)
    """
    roi_type = roi.getType()
    poly = roi.getFloatPolygon()
    roi_name = roi.getName() if roi.getName() else ""


    if roi_type in (Roi.POLYGON, Roi.FREEROI, Roi.TRACED_ROI, Roi.POLYLINE,
Roi.FREELINE, Roi.ANGLE):
        # general polygon-based approach
        new_x = [(x * scale_x + translate_x) for x in poly.xpoints]
        new_y = [(y * scale_y + translate_y) for y in poly.ypoints]
```

```
        new_roi = PolygonRoi(new_x, new_y, len(new_x), roi_type)

    elif roi_type == Roi.RECTANGLE:
        # transform bounding rectangle
        bounds = roi.getBounds()
        new_x = bounds.x * scale_x + translate_x
        new_y = bounds.y * scale_y + translate_y
        new_w = bounds.width * scale_x
        new_h = bounds.height * scale_y
        new_roi = Roi(new_x, new_y, new_w, new_h)

    elif roi_type == Roi.LINE:
        # transform endpoints
        if len(poly.xpoints) == 2:
            x1 = poly.xpoints[0] * scale_x + translate_x
            y1 = poly.ypoints[0] * scale_y + translate_y
            x2 = poly.xpoints[1] * scale_x + translate_x
            y2 = poly.ypoints[1] * scale_y + translate_y
            new_roi = Line(x1, y1, x2, y2)
        else:
            # Possibly skip or handle differently
            return None
    else:
        # unsupported type => skip
        return None

    new_roi.setName(roi_name)
    return new_roi

def transform_rois(rois, ref_roi, user_roi):
    """
```

```
    Takes pattern rois, the reference bounding box ROI, and the user bounding box ROI.
    Returns the newly transformed ROIs.
    """
    ref_bounds = ref_roi.getBounds()
    user_bounds = user_roi.getBounds()

    # Compute scaling + translation
    scale_x = user_bounds.width / float(ref_bounds.width)
    scale_y = user_bounds.height / float(ref_bounds.height)
    translate_x = user_bounds.x - ref_bounds.x * scale_x
    translate_y = user_bounds.y - ref_bounds.y * scale_y

    transformed = []
    for roi in rois:
        new_roi = transform_single_roi(roi, scale_x, scale_y, translate_x, translate_y)
        if new_roi:
            transformed.append(new_roi)

    return transformed

def compute_rotation_angle(line_roi):
    """
    Computes rotation angle from a 2-point line ROI,
    returning an angle that avoids flipping the image upside down.
    """
    poly = line_roi.getFloatPolygon()
    x1, y1 = poly.xpoints[0], poly.ypoints[0]
    x2, y2 = poly.xpoints[1], poly.ypoints[1]
    angle = math.degrees(math.atan2(y2 - y1, x2 - x1))

    rotate_angle = -angle
```

```
    # Prevent over-rotation
    if rotate_angle > 90:
        rotate_angle -= 180
    elif rotate_angle < -90:
        rotate_angle += 180

    return rotate_angle

# ========================
# Main Script
# ========================

# Ensure ROI Manager Closed at Start
rm_existing = RoiManager.getInstance()
if rm_existing:
    rm_existing.close()

# Prompt for working directory containing converted .tiff s
dc = DirectoryChooser("Choose the Converted .tiff Directory")
work_dir = dc.getDirectory()

# Create subfolders if not present
output_dirs = ["Channel1", "Channel2", "Merged", "ROI"]
for odir in output_dirs:
    dir_path = os.path.join(work_dir, odir)
    if not os.path.exists(dir_path):
        os.makedirs(dir_path)

# Prompt for ROI pattern ZIP
roi_zip_path = OpenDialog("Choose ROI Pattern ZIP file").getPath()
rm_pattern = RoiManager()
```

```
rm_pattern.reset()
rm_pattern.runCommand("Open", roi_zip_path)

pattern_rois = [rm_pattern.getRoi(i) for i in range(rm_pattern.getCount())]
ref_bbox_roi = next((r for r in pattern_rois if r.getName() == "BoundingBox"), None)
if ref_bbox_roi is None:
    IJ.error("BoundingBox ROI not found. Exiting.")
    exit()

# Clear ROI Manager after verifying bounding box
rm_pattern.reset()

# ========================
# Process each TIF file
# ========================
for file_name in os.listdir(work_dir):
    if not file_name.lower().endswith(".tif"):
        continue

    file_path = os.path.join(work_dir, file_name)
    imp = IJ.openImage(file_path)
    if not imp:
        IJ.log("Failed to open: " + file_name)
        continue

    # Split into channels
    channels = ChannelSplitter.split(imp)
    ch1, ch2 = channels[0], channels[1]

    ch1.show()  # channel 1 is brightfield / interactive
    IJ.setTool("line")
```

```
# Initialize new ROI Manager for deskew
rm = RoiManager.getInstance()
if rm is None:
    rm = RoiManager()
rm.reset()

# Prompt for deskew line
WaitForUserDialog("Deskewing",
    "Draw a line along feature on channel 1. Add the line to ROI Manager, then click OK."
).show()

if rm.getCount() < 1 or rm.getRoi(0).getType() != Roi.LINE:
    IJ.log("Invalid deskew line. Skipping file: " + file_name)
    ch1.close()
    continue

# Compute & apply rotation
angle = compute_rotation_angle(rm.getRoi(0))
IJ.run(ch1, "Rotate...", "angle={} interpolation=Bilinear enlarge".format(angle))
IJ.run(ch2, "Rotate...", "angle={} interpolation=Bilinear enlarge".format(angle))
rm.reset()

# Prompt for bounding box
IJ.setTool("rectangle")
WaitForUserDialog("ROI Alignment",
    "Draw a rectangular bounding box on channel 1. Add it to ROI Manager, then click OK."
).show()

if rm.getCount() < 1 or rm.getRoi(0).getType() != Roi.RECTANGLE:
    IJ.log("Invalid bounding box. Skipping file: " + file_name)
```

```
        ch1.close()
        ch2.close()
        continue

    user_bbox_roi = rm.getRoi(0)
    transformed = transform_rois(pattern_rois, ref_bbox_roi, user_bbox_roi)

    # Save the newly transformed ROIs
    rm.reset()
    for r in transformed:
        rm.addRoi(r)

    roi_save_path = os.path.join(work_dir, "ROI", file_name.replace(".tif", "_ROIs.zip"))
    rm.runCommand("Save", roi_save_path)

    # Save processed channels
    FileSaver(ch1).saveAstiff(os.path.join(work_dir, "Channel1", file_name))
    FileSaver(ch2).saveAstiff(os.path.join(work_dir, "Channel2", file_name))

    # Merge & save
    merged = RGBStackMerge.mergeChannels([ch1, ch2], False)
    FileSaver(merged).saveAsTiff(os.path.join(work_dir, "Merged", file_name))

    # Cleanup
    ch1.close()
    ch2.close()
    merged.close()
    rm.reset()

IJ.showMessage("Processing completed successfully!")
```

***Supplementary Macro 3: Condensation Quantification***

```
# Fluorescence Analysis in Fiji using Jython (Python)
from ij import IJ
from ij.io import DirectoryChooser
from ij.plugin.frame import RoiManager
from ij.plugin import Duplicator
from ij.measure import Measurements
from java.io import FileWriter, BufferedWriter
import os

# --- Prompt for folders ---
imgDir = DirectoryChooser("Select folder containing .tiff files").getDirectory()
roiDir = DirectoryChooser("Select folder containing ROI ZIPs").getDirectory()
outputFile = os.path.join(imgDir, "Fluorescence.csv")
maskOutputDir = os.path.join(imgDir, "Thresholded_Masks")

if not os.path.exists(maskOutputDir):
    os.makedirs(maskOutputDir)

# --- Create CSV output ---
writer = BufferedWriter(FileWriter(outputFile))
writer.write("Image,ROI,Signal_Area_um2,Total_Area_um2,%_Signal_Coverage\n")

# --- Process each .tiff image ---
for fileName in os.listdir(imgDir):
    if not (fileName.lower().endswith(".tif") or fileName.lower().endswith(".tiff")):
        continue

    imgPath = os.path.join(imgDir, fileName)
    imp = IJ.openImage(imgPath)
    if imp is None:
```

```
        print("Could not open image:", imgPath)
        continue

    baseName = os.path.splitext(fileName)[0]
    roiZip = os.path.join(roiDir, baseName + "_ROIs.zip")
    if not os.path.exists(roiZip):
        print("ROI ZIP not found:", roiZip)
        imp.close()
        continue

    # --- Create thresholded binary mask ---
    imp_thresh = Duplicator().run(imp)
    IJ.run(imp_thresh, "8-bit", "")
    IJ.setAutoThreshold(imp_thresh, "RenyiEntropy")
    IJ.run(imp_thresh, "Make Binary", "")
    IJ.run(imp_thresh, "Convert to Mask", "")  # Lock into 0/255

    # --- Save thresholded image ---
    thresholdOutputPath = os.path.join(maskOutputDir, "Thresh_" + baseName + ".tif")
    IJ.saveAsTiff(imp_thresh, thresholdOutputPath)

    # --- Load ROIs ---
    rm = RoiManager.getRoiManager()
    rm.reset()
    rm.runCommand("Open", roiZip)
    rois = rm.getRoisAsArray()
    if not rois or len(rois) == 0:
        print("No ROIs found in:", roiZip)
        imp_thresh.close()
        imp.close()
        continue
```

```
    # --- Calibration for µm² ---
    cal = imp.getCalibration()
    pixelArea = cal.pixelWidth * cal.pixelHeight

    for roi in rois:
        roiName = roi.getName()

        # Total ROI area from original image
        imp.setRoi(roi)
        totalStats = imp.getStatistics(Measurements.AREA)
        totalArea = totalStats.area * pixelArea

        # Signal area (white pixels = 255 in mask)
        imp_thresh.setRoi(roi)
        stats = imp_thresh.getStatistics()
        hist = stats.getHistogram()
        signalPixels = hist[255]
        signalArea = signalPixels * pixelArea

        # % signal coverage
        percentCoverage = (signalArea / totalArea) * 100 if totalArea > 0 else 0

        row = "{},{},{:.3f},{:.3f},{:.2f}\n".format(fileName, roiName, signalArea, totalArea,
percentCoverage)
        writer.write(row)

    rm.reset()
    imp_thresh.close()
    imp.close()
```

```python
writer.close()

# --- Post-step: Invert all saved masks (overwrite in place) ---
for fileName in os.listdir(maskOutputDir):
    if not fileName.lower().endswith(".tif"):
        continue

    filePath = os.path.join(maskOutputDir, fileName)
    imp_mask = IJ.openImage(filePath)
    if imp_mask is None:
        print("Could not open mask:", filePath)
        continue

    ip_mask = imp_mask.getProcessor()
    ip_mask.invert()
    imp_mask.updateAndDraw()

    IJ.saveAsTiff(imp_mask, filePath)
    imp_mask.close()

IJ.showMessage("Done", "Fluorescence.csv saved and all masks inverted in place.")
```
